# Accelerated, Accurate, Hybrid Short and Long Reads Alignment and Variant Calling

**DOI:** 10.1101/2025.04.15.648987

**Authors:** Jinnan Hu, Don Freed, Hanying Feng, Hong Chen, Zhipan Li, Haodong Chen

**Affiliations:** Sentieon Inc., San Jose, CA, USA

## Abstract

**Background:** Integrating short-read and long-read sequencing technologies has become a promising approach for achieving accurate and comprehensive genomic analysis. While short-read sequencing (Illumina, etc.) offers high base accuracy and cost efficiency, it struggles with structural variation (SV) detection and complex genomic regions. In contrast, long-read sequencing (PacBio HiFi) excels in resolving large SVs and repetitive sequences but is limited by throughput, higher Indel error rates, and sequencing costs. Hybrid approaches may combine these technologies and leverage their complementary strengths and different sources of error to provide higher accuracy, more comprehensive results, and higher throughput by lowering the coverage requirement for the long reads.

**Methods:** This study benchmarks the DNAscope Hybrid pipeline, a novel integrated alignment and variant calling framework that combines short- and long-read data sequenced from the same sample. We evaluate its performance across multiple human genome reference datasets (HG002–HG004) using the draft Q100 and Genome in a Bottle v4.2.1 benchmarks. The pipeline’s ability to detect small variants (SNPs/Indels), structural variants (SVs), and copy number variations (CNVs) is assessed using data from the Illumina and Pacbio sequencing systems at varying read depths (5x–30x). Benchmark results are compared to DeepVariant.

**Results:** The DNAscope Hybrid pipeline significantly improves SNP and Indel calling accuracy, particularly in complex genomic regions. At lower long-read depths (e.g., 5x-10x), the hybrid approach outperforms standalone short- or long-read pipelines at full sequencing depths (30x-35x). Additionally, the DNAscope Hybrid outperforms leading open-source tools for SV and CNV detection, enhancing variant discovery in challenging genomic regions. The pipeline also demonstrates clinical utility by identifying disease-associated variants. Moreover, DNAscope Hybrid is highly efficient, achieving less than 90 minutes runtimes at single standard instance.

**Conclusion:** The DNAscope Hybrid pipeline is a computationally efficient, highly accurate variant calling framework that leverages the advantages of both short- and long-read sequencing. By improving variant detection in challenging genomic regions and offering a robust solution for clinical and large-scale genomic applications, it holds significant promise for genetic disease diagnostics, population-scale studies, and personalized medicine.

## Introduction

Over the past decade, next-generation sequencing (NGS) and third-generation sequencing (TGS) have become a cornerstone in genomics research and medical applications, driving significant discoveries in disease mechanisms, population diversity, and personalized medicine strategies^1,2^. These advancements were facilitated by improvements in sequencing technologies, including reduced costs, enhanced read lengths, higher base quality, and increased accessibility to laboratories at various sizes.

Highly accurate methods for detecting single-nucleotide polymorphisms (SNPs) and <50bp insertions or deletions (Indels) have been central to genetic disease and tumor diagnostics. Additionally, the adoption of long-read sequencing has enabled better integration of structural variation (SV, ≥50 bp) into analyses^3,4^. Although SVs are less abundant than small variants in the human genome, they collectively impact more base pairs and play crucial roles in human evolution and disease^5^. Copy number variations (CNVs), arising from DNA segment deletions or duplications, represent another form of genomic variation linked to various diseases^6^. Despite these advancements, detecting and interpreting these variants together in an integrated analysis pipeline remain challenging.

While short-read sequencing technologies (e.g., Illumina, Element Biosciences, MGI, etc.) effectively capture small variations across most of the human genome, they face challenges in difficult-to-map regions and in the detection of structural variation (SV). Studies have demonstrated the limitations of short reads for identifying larger insertions, deletions, and other complex genomic rearrangements^7^. Long-read sequencing technologies, such as PacBio HiFi and others, have been proposed to address these issues. These platforms enable improved detection of complex SVs due to their ability to produce reads exceeding 15 kb in length with current base accuracies ranging from 99% to 99.9%^3,8,9^. Nevertheless, these technologies are not without challenges. Errors in long-read sequencing often manifest as context-specific insertions and deletions (e.g., homopolymers), complicating the detection of indel variants even with high read coverage^10^. Additionally, the high cost of generating long reads, combined with their computational demands, poses barriers to large-scale applications, including population-wide studies and analysis of legacy samples. Many interesting samples slated for long read analysis, already have ∼30x short read data. By using 30x short read data with long read data, this new pipeline leverages the strengths of both technologies, allowing us to lower the typical long read coverage needed by 2-3x while simultaneously increasing the accuracy and comprehensiveness of results for each individual sample.

The complementary error profiles of short- and long-read sequencing technologies have motivated the development of hybrid analysis pipelines that leverage both data types. Initially, such approaches were implemented for de novo genome assembly, where short reads were used to correct errors in long-read assemblies^11,12^. Several hybrid re-sequencing pipelines have also emerged, including “HELLO,” which employs deep learning to perform variant calling using combined alignments of short and long reads^13^. Another notable pipeline, “blend-seq,” focuses on combining ultra-low coverage long reads (∼4×) with standard 30× short reads for cost-effective variant discovery^14^. Clinically, Variantyx has integrated short- and long-read analyses into a single diagnostic workflow, generating a comprehensive clinical report. This pipeline, however, uses long reads primarily for orthogonal confirmation of variants detected by short reads, leaving opportunities for further integration and optimization^15^.

These existing pipelines independently align short and long reads to reference genomes without exploiting the potential of realignment to add value for variant calling. Moreover, limited attention is given to computational efficiency and speed, making them less viable for clinical settings like neonatal intensive care units (NICUs) or large-scale cohort analyses.

The Genome in a Bottle Consortium (GIAB) has progressively improved its reference sample variant benchmark. The v4.2.1 variant call set, released in 2022, incorporated linked-reads and long-read sequencing data, expanding high-confidence regions in the GRCh38 assembly from 85% to 92% of the genome. This update introduced difficult-to-map regions and other challenging genomic loci not previously included in the v3.3.2 call set^16^. In addition to genome-wide SNP/indel benchmark, the GIAB released an SV benchmark (v0.6)^7^ and a benchmark for Challenging Medically Relevant Genes (CMRG)^17^. In parallel, the Telomere-to-Telomere (T2T) Consortium has published high-quality assemblies of the HG002 sample^18^. The initial assembly leveraged PacBio HiFi and ONT data from the Human Pangenome Reference Consortium (HPRC) and GIAB. Following extensive polishing and validation, the v1.1 diploid assembly achieved near-perfect haplotype phasing and an error rate below one per 10 billion bases (a Phred quality score of Q100)^19^. Through alignment of the Q100 assembly to GRCh38, the GIAB team has generated a draft assembly-based benchmark for HG002. This new benchmark provides significantly more small variants and nearly three times the number of confident SV events compared to the earlier GIAB v0.6 SV benchmark (30,244 vs. 9,646)^20^. These advancements underscore the importance of choosing technologies and datasets aligned with cutting-edge genomic knowledge for clinical and research applications.

Sentieon has won a variety of awards in the in PrecisionFDA Challenges including an award in Truth Challenge V2 for multi-platform analysis,^21^ where short and long-reads were used to improve accuracy. The DNAscope Hybrid pipeline presented here is a substantial improvement from the PrecisionFDA winning pipeline. Different from the previously published DNAscope pipeline for short-reads^22^, and the DNAscope LongRead pipeline for long-reads^23^, this hybrid analysis tool integrates short- and long-read sequencing data from the same sample to deliver comprehensive and accurate variant calling.

As shown in Figure 1, DNAscope Hybrid accepts Fastq or BAM files as input and generates VCF outputs containing SNP, indel, SV, and CNV data. By combining the strengths of both sequencing platforms, the pipeline achieves superior variant detection compared to using either technology in isolation. DNAscope Hybrid can be used with whole-genome sequencing (WGS) long-read data or with targeted sequencing approaches like the Twist Alliance Dark Genes Panel^24^. DNAscope Hybrid’s performance and versatility make it a promising tool for clinical diagnostics, particularly in settings requiring highly accurate, comprehensive results.

**Figure 1:**
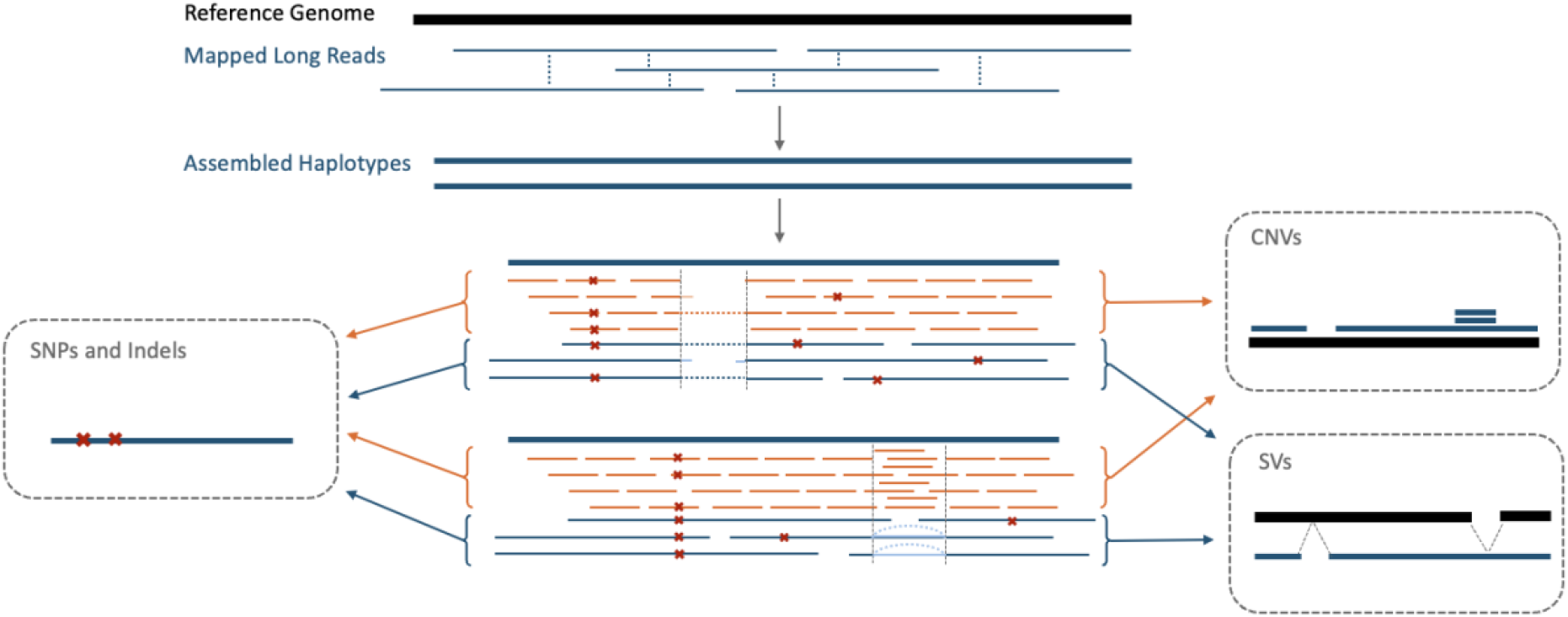
Overview processing steps of DNAscope Hybrid variant calling pipeline.

## Results

### Small Variants (SNPs and Indels)

To evaluate the accuracy of the DNAscope Hybrid (DS-Hybrid) pipeline at varying depths, we used the HG002 sample and down sampled PacBio HiFi (PB) datasets to depths of 5x, 7.5x, 10x, 15x, 20x, and 30x, pairing them with 35x Illumina (ILMN) short-read data. To assess the accuracy contribution of short reads, we also analyzed each depth of PB datasets independently without short reads using the DNAscope LongRead (DS-LR) pipeline. Additionally, we included other datasets for comparison: ILMN (35x) analyzed with Dragen v4.2, and PacBio HiFi (30x) analyzed by DeepVariant (v1.8.0).

We initially investigated genome-wide accuracy using the NIST v4.2.1 benchmark (Figure 2 A-B, Supplementary Table S1). This analysis demonstrated that higher depths of long-read sequencing yield greater accuracy, with the highest accuracy observed in the combined 30x PB + 35x ILMN datasets. Further, Hybrid Indel accuracy is much higher than all other methods, even when using only 7.5x coverage for long reads. Notably, the DNAscope Hybrid pipeline dramatically improved SNP and Indel accuracy compared to any single-technology pipeline.

**Figure 2.**
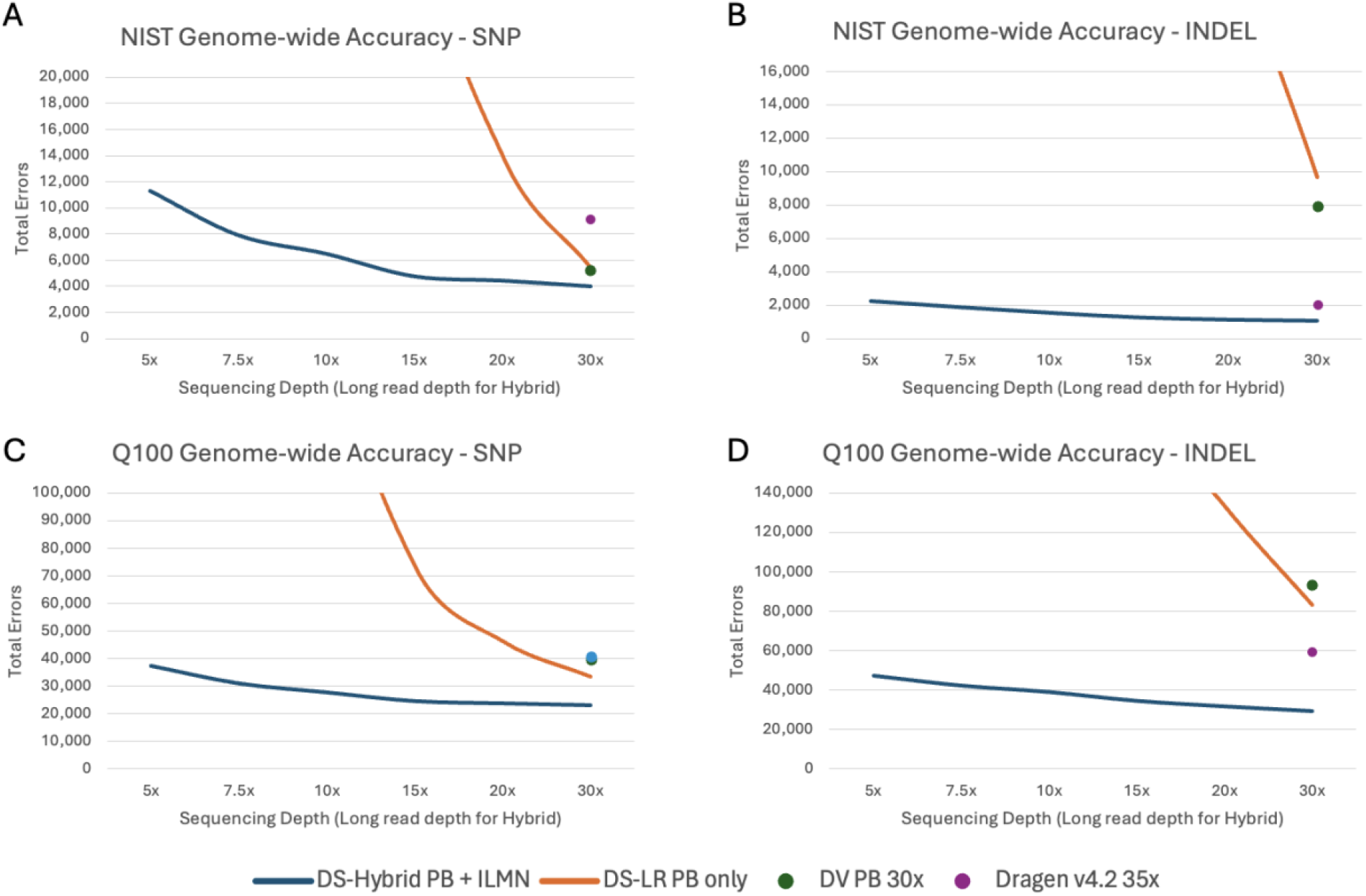
Genome-wide Accuracy - Total Errors of A) SNP in GIAB v4.2.1; B) Indel in GIAB v4.2.1; C) SNP in draft Q100; D) Indel in draft Q100. DS-Hybrid PB+ILMN and DS-LR PB only are shown with curves covering 5x-30x long reads depths. DV PB and Dragen are shown at full depth.

The current cost for DNA extraction and library preparation is approximately $735 USD for PacBio HiFi and $135 USD for Illumina (the service cost is from a single service provider^25^, as it will differ elsewhere). Sequencing costs are around $330 for 10x PacBio HiFi coverage^26^ and $200 for 30x Illumina coverage^27^. Therefore, generating a combined dataset of 10x PacBio HiFi plus 30x Illumina would result in a total wet lab cost comparable to generating 20x PacBio HiFi data alone.

Based on this data, 10x of Pacbio and 35x of Illumina has a good blend of cost of reagents vs results. At this coverage level the pipeline gets only 1527 Indel errors and only 6467 SNP errors for an F1 of 0.9985 and 0.9990 respectively.

Comparing the draft Q100 and the v4.2.1 benchmarks, total errors are much higher with the draft Q100 benchmark as it contains more challenging regions (Figure 2 C-D, Table S2), making it more suitable for benchmarking new high accuracy variant callers. In the draft Q100 benchmark, the hybrid pipeline has fewer errors relative to single-technology pipelines for both SNPs and Indels. Comparing the hybrid pipeline with 10x long-read coverage with the draft Q100, SNP errors are reduced by 30% relative to the next-best pipeline (DS-LR) and Indel errors are reduced by 35% relative to the next-best pipeline (Dragen).

To better understand the variant calling accuracy improvement in the hybrid pipeline, we performed a stratified analysis across GA4GH stratification regions^28^. Variant calling accuracy as measured with the draft Q100 benchmark at annotated tandem repeats and homopolymer (TRHP) regions is shown in Figure 3 A-B. We additionally assessed variant calling accuracy using the CMRG benchmark for HG002 (Figure 3 C-D). The DNAscope Hybrid pipeline has improved accuracy at regions, as even 5x Hybrid call sets outperforms any other high depth call sets, for both SNPs and Indels. Short reads frequently fail to map to tandem repeats correctly, and long reads have less accurate resolution of homopolymers – by using the two datatypes in a complimentary way, the Hybrid method helps resolve both sources of error. CMRG regions, which encompass 273 medically relevant genes, demonstrated substantial benefits from hybrid short and long-read data. Long reads alone cannot capture each variant correctly, while the Hybrid pipeline still showed its improved accuracy, especially for Indels. The improved accuracy will likely lead to an improved diagnostic rate and other clinical utility.

**Figure 3.**
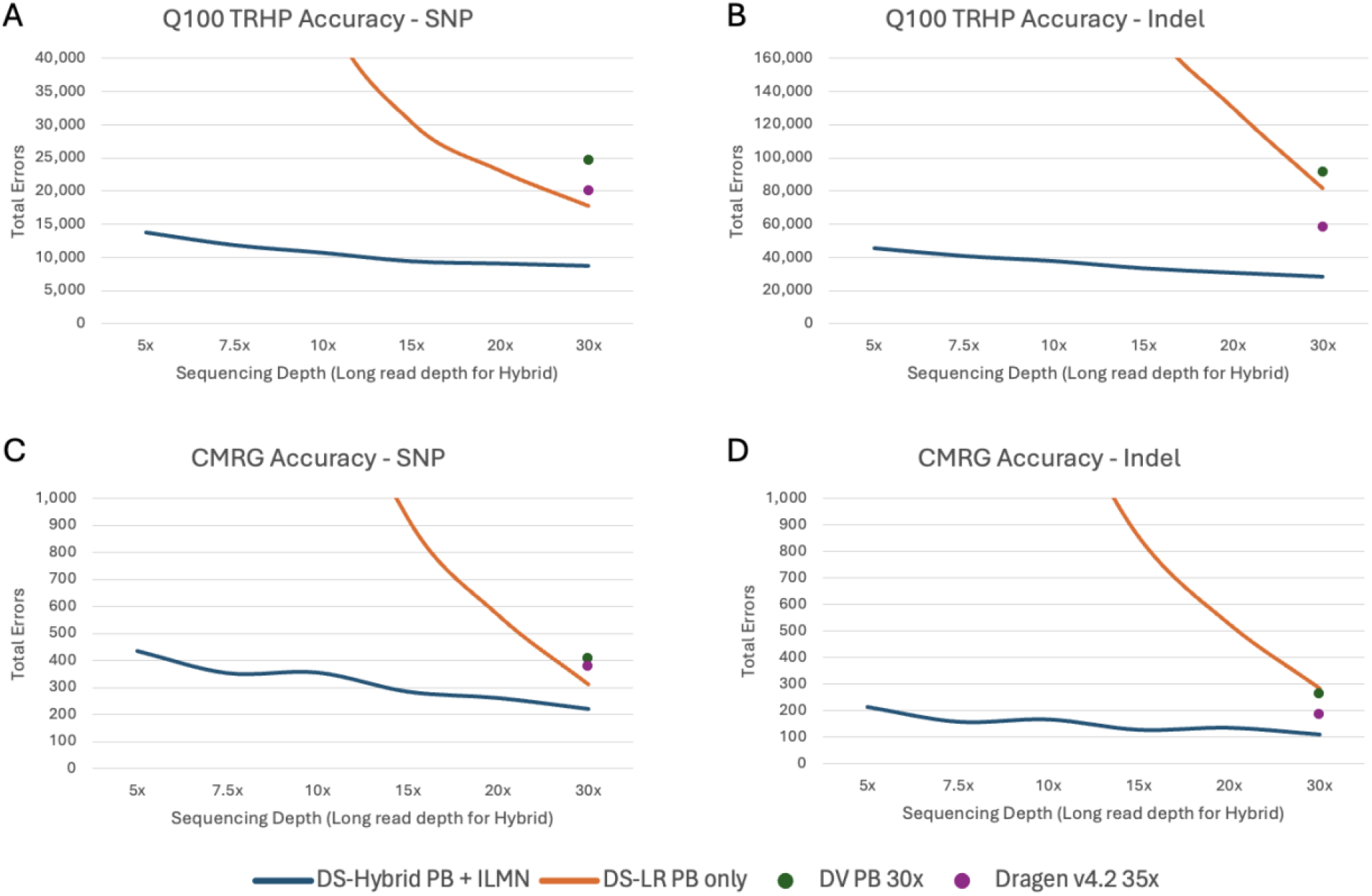
Stratified Region Accuracy - Total Errors of A) SNP and B) INDEL in tandem repeat and homopolymer regions (TRHP) Q100 benchmark. C) SNP and D) Indel in challenging medically relevant gene (CMRG) regions. DS-Hybrid PB+ILMN and DS-LR PB only are shown with curves covering 5x-30x long reads depths. DV PB and Dragen are shown at full depth.

To better understand the differences between the evaluated pipelines, we compared the intersection of the Hybrid and standalone short or long-read pipelines together with the benchmark VCF. Variants detected by all pipelines and also present in the benchmark VCF represent the highest proportion but are not displayed in the figures (Figure 4). ILMN detected fewer SNPs, with a higher number of false negatives, while PB had a higher rate of false positive SNPs. Variants missed by short-read pipelines were mainly attributed to low mappability and poor coverage, while those missed by long-read pipelines were primarily due to inherent limitations in base-calling accuracy, particularly at homopolymer indels.

**Figure 4.**
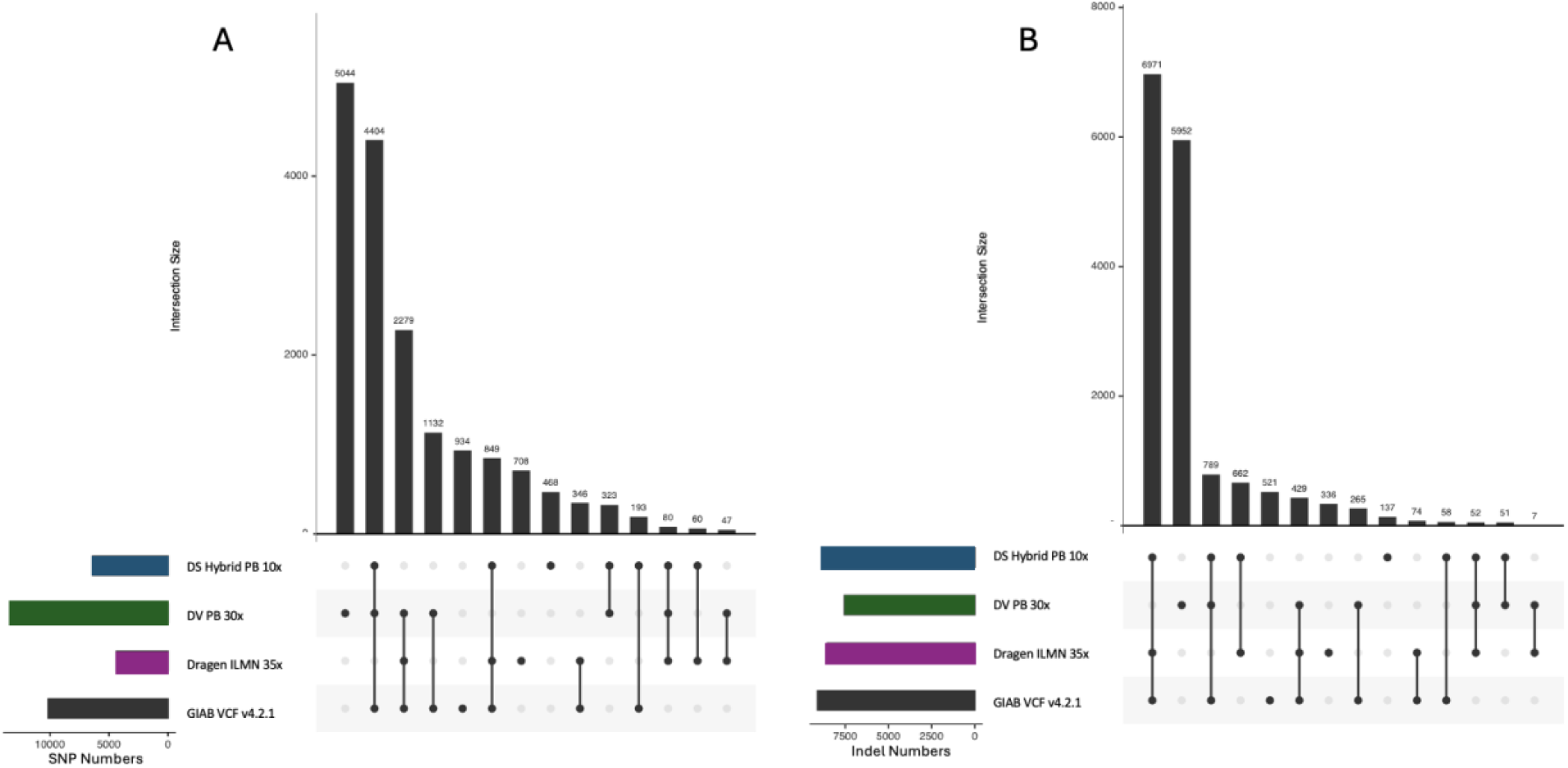
UpSet plots of (A) SNPs and (B) Indels, including 3 benchmarked pipelines and the GIAB v4.2.1 benchmark VCF. The intersection (bar) of variants identified by all 3 pipelines that are also in the GIAB VCF is removed, and the remaining intersection categories are sorted by size. Call set sizes for each pipeline are displayed on the lower left panels. The UpSet plots have slightly different results from that in Figure 2 and Table S1, due to different comparison approached used.

While the DNAscope Hybrid pipeline has excellent performance on HG002, we wanted to further assess the performance on additional datasets to ensure that the approach used by the pipeline extends to other samples. We then applied the DNAscope Hybrid pipeline to two additional GIAB samples. The results, measured as SNP and Indel combined total errors (FP+FN), were compared to the platform recommended pipelines for Illumina short reads and PacBio Long Reads (Figure 5). Figure 5 shows DNAscope Hybrid call sets are consistently more accurate than short or long reads only call sets in every sample, highlighting its robustness and adaptability.

**Figure 5.**
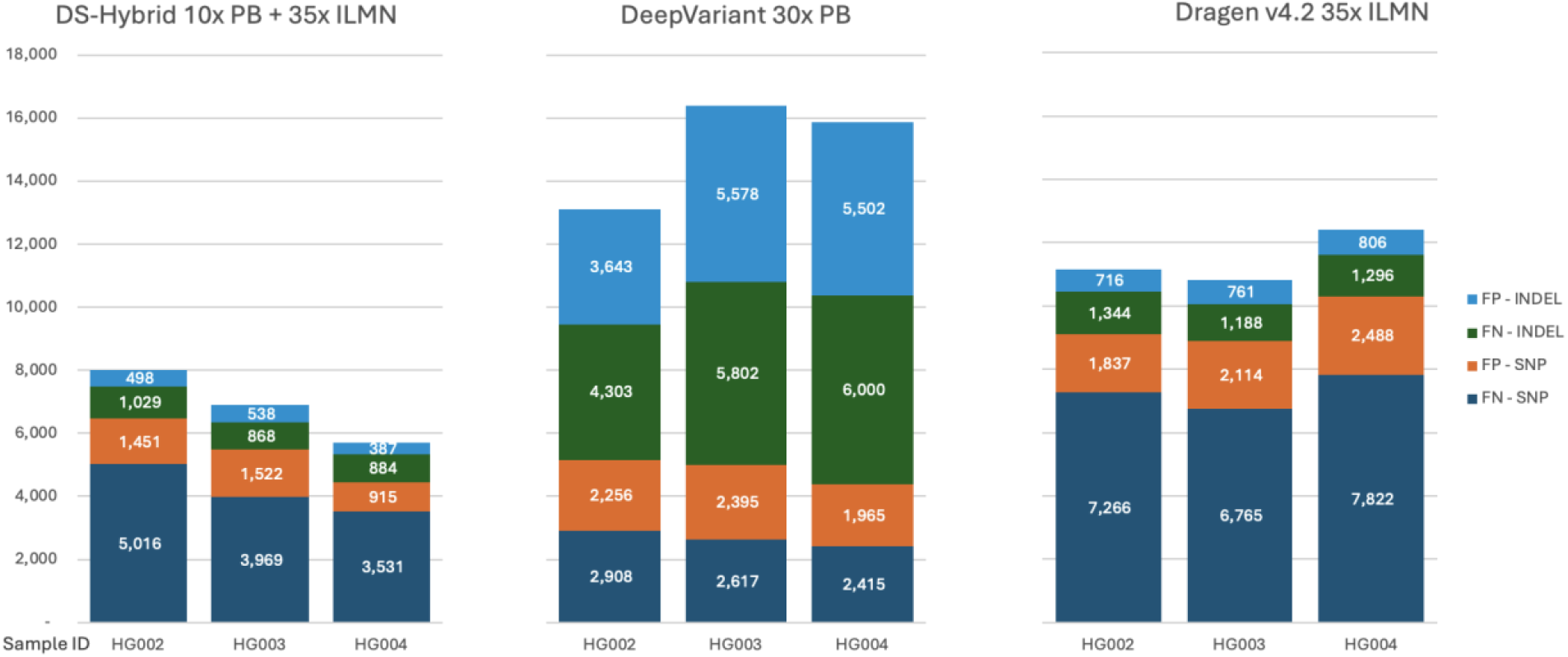
SNP/Indel accuracy over HG002-004 reference samples. False positive and false negative of SNP and Indel are listed separately.

### Structural Variants (SVs)

To evaluate structural variant (SV) accuracy, we analyzed down sampled long- and short-read datasets, and evaluated variant calling accuracy using the draft Q100 or the CMRG SV benchmark. For the hybrid SV pipeline, only long-read information was utilized, so the accuracy represents both DNAscope Hybrid and DNAscope LongRead. Other benchmarked pipelines include PacBio SV calls generated using Pbsv and 35x ILMN SV calls generated using Dragen v4.2.

The genome-wide draft Q100 SV benchmark complements the information in the SNP/Indel accuracy curves (Figure 6 A-B, Table S6). While short-read data pipelines demonstrate high accuracy for SNP and Indel detection, these pipelines struggled at SV calling, especially recall. In contrast, long-read data shows much higher SV accuracy, even at lower depths, and achieved saturation in performance at a depth around 15-20x. Notably, the DNAscope Hybrid/LR pipeline outperformed Pbsv in this benchmark. To further validate performance, we assessed SV accuracy using the CMRG SV benchmark (Figure 6 C-D, Table S7). Performance on the CMRG SV benchmark was consistent with the performance observed in the larger draft Q100 SV benchmark, underscoring the advantage of long-read sequencing in SV detection.

**Figure 6.**
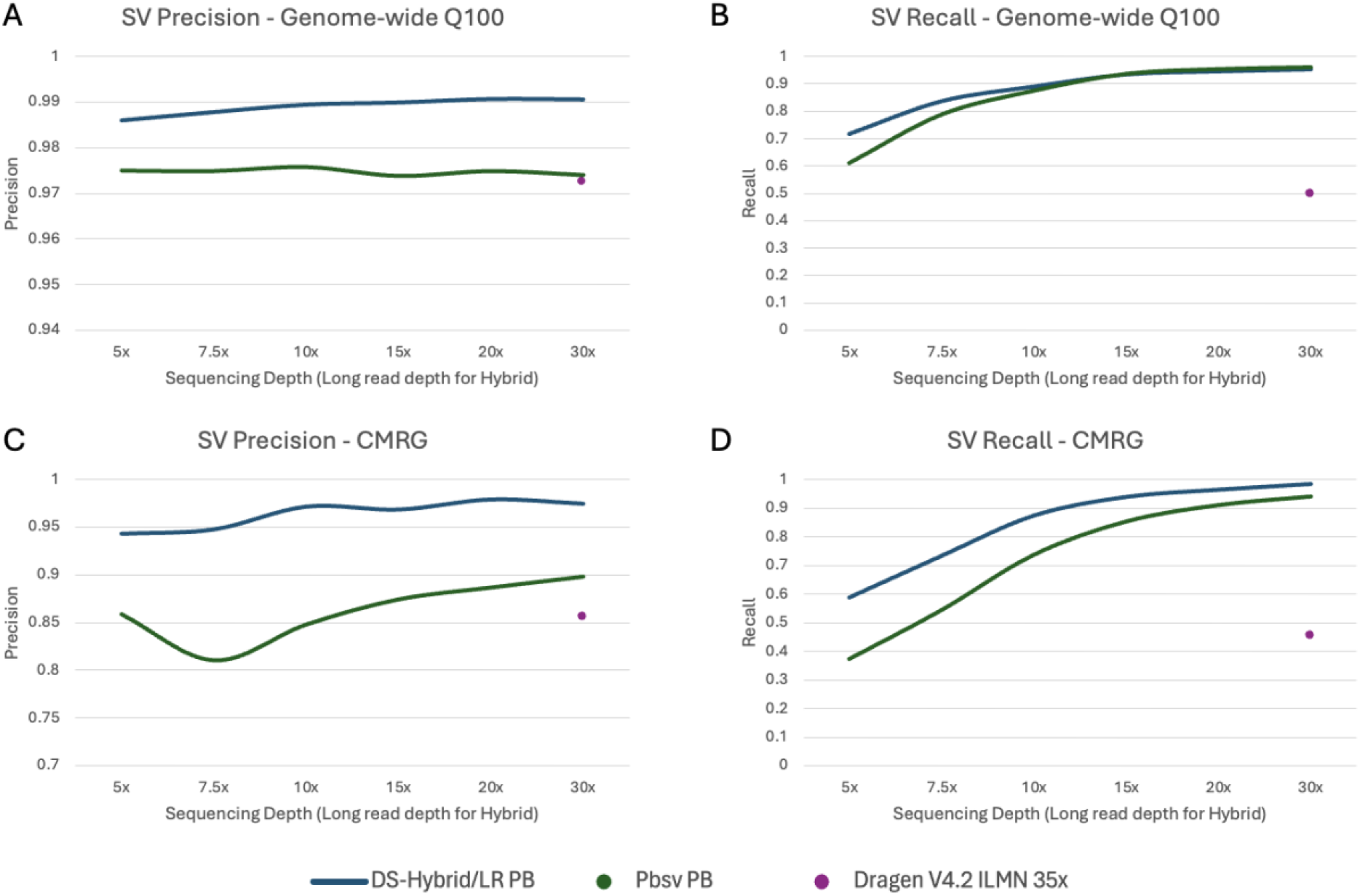
Structural variation (SV) accuracy as measured by A) precision on the draft Q100 benchmark; B) recall on the draft Q100 benchmark; C) Precision on the CMRG SV benchmark; D) Recall on the CMRG SV benchmark. PB-Hybrid/LongRead and Pbsv PB are shown as curves covering 5x-30x long-read depths. Dragen ILMN is showing at full depth accuracy.

We further analyzed the intersection of the SV call sets with the draft Q100 SV benchmark (Figure 7). This analysis highlights the substantial number of structural variants missed by the short-read pipeline. These omissions primarily stem from the inherent limitations of short reads, particularly their inability to span the long sequences characteristic of many SVs.

**Figure 7.**
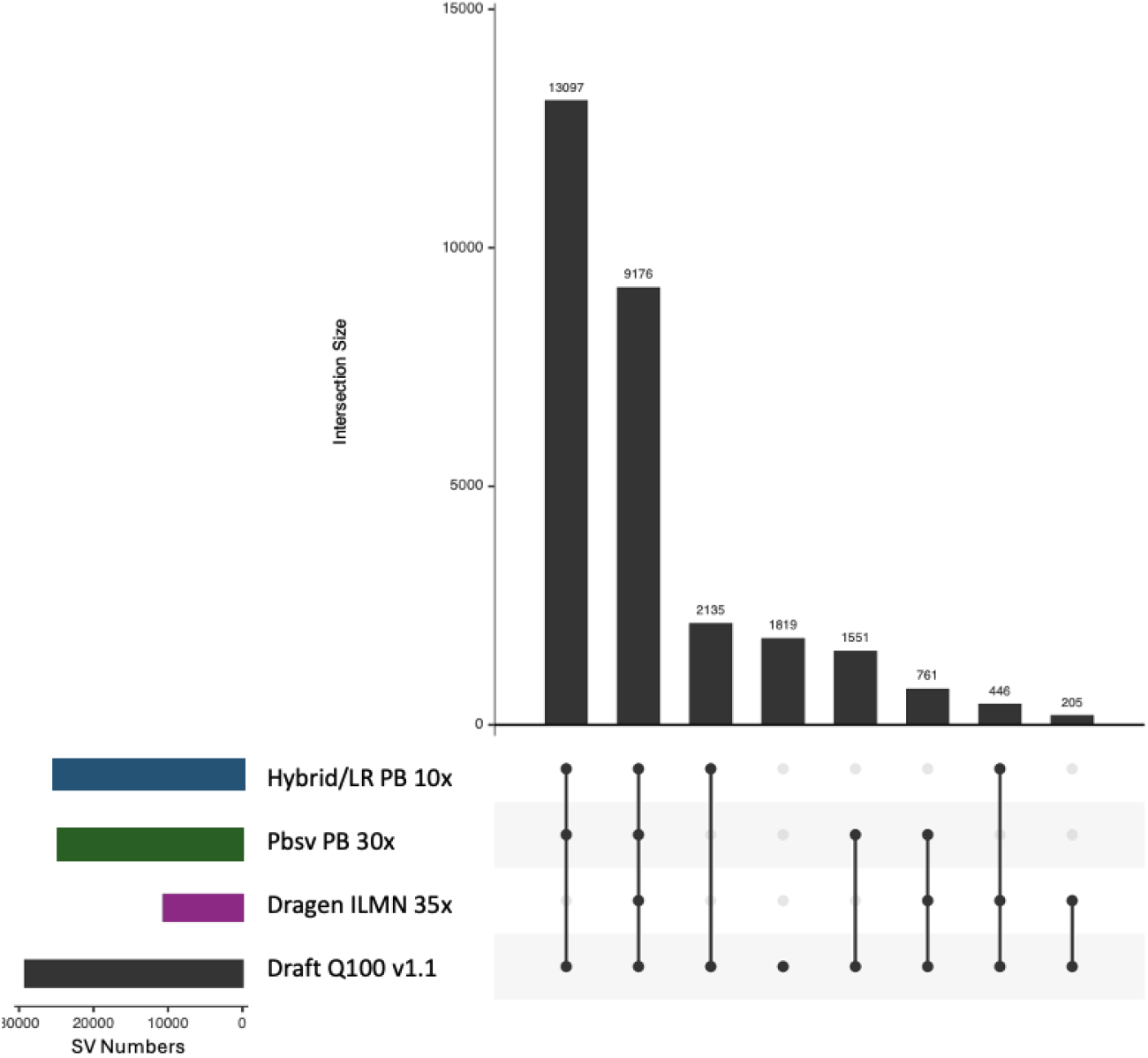
Upset figures of SVs, including 3 benchmarked pipelines and the draft Q100 benchmark. Intersection categories are sorted by size. Call set sizes for each pipeline are displayed on the lower left panels.

### Copy Number Variation (CNV)

CNV refers to genetic differences between individuals involving the loss or gain of specific DNA regions. The current version of the DNAscope Hybrid pipeline utilizes the recently released Sentieon CNVscope module, which relies solely on short-read data for CNV detection. To evaluate its performance, we benchmarked CNVscope accuracy using the HG002 Q100 benchmark (Figure 8; see methods). We also benchmarked CNVnator accuracy across different size lengths.

**Figure 8.**
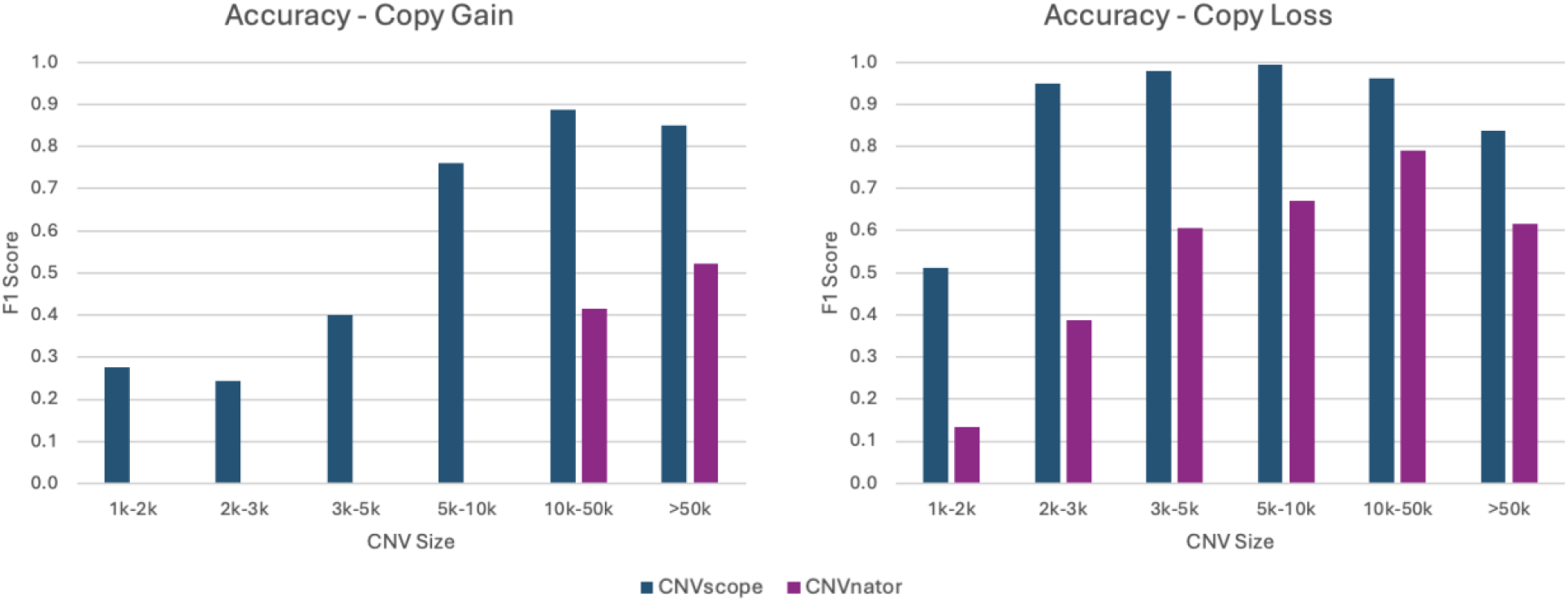
CNV accuracy benchmark. CNVscope serves as the CNV caller in the DNAscope Hybrid and short-reads pipelines. CNVnator results are shown for comparison.

The Sentieon pipeline consistently demonstrated higher accuracy across nearly all event sizes, including the technically challenging <10k events, where CNVnator and most other tools struggle to achieve high accuracy. This suggests that DNAscope Hybrid offers significant improvements in CNV detection.

### Overall Variant Counts in HG002

DNAscope Hybrid is a comprehensive variant-calling pipeline capable of detecting SNVs, short Indels (<50 bp) and longer insertions and deletions. When compared with the short reads Dragen pipeline, The Hybrid pipeline identified a notably higher number of variants, particularly large Insertion events (Figure 9). This increase is primarily attributable to the additional information provided by long-read data integrated into the hybrid workflow.

**Figure 9.**
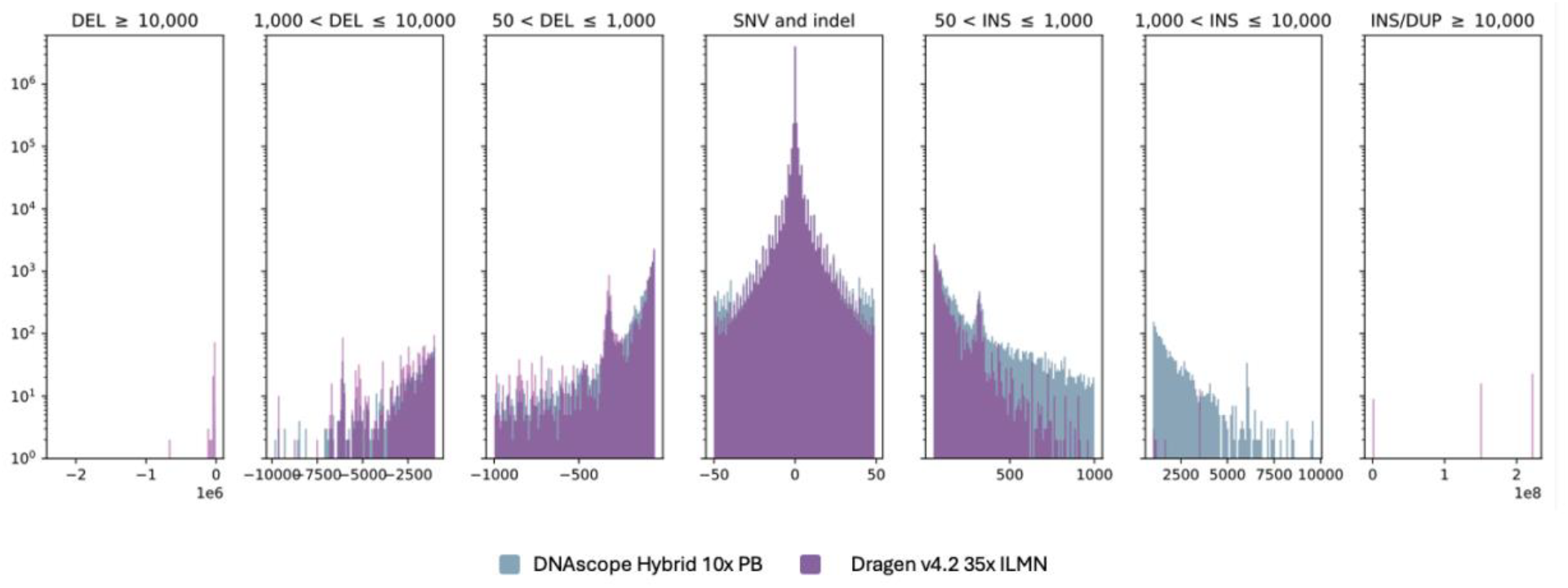
**Size distribution of small and structural variants** identified by DNAscope Hybrid pipeline on 10x PB + 35x ILMN, and by Dragen on 35x ILMN.

### Clinical Validation on simulated pathogenic variants

To evaluate the clinical utility of the DNAscope Hybrid pipeline, we further analyzed HG002 variants in the CMRG benchmark. Specifically, HG002 CMRG variants were annotated using VEP^29^, and those associated with exons were selected for comparison across three pipelines: 1) the DNAscope Hybrid with 10x PB and 35x ILMN; 2) DeepVariant 30x PB; 3) DeepVariant 35x ILMN. Variants detected by only one or two of these pipelines are shown in the landscape figure (Figure 10). The DNAscope Hybrid pipeline identified all 64 variants, whereas DV PB and DV ILMN failed to capture some, with nine variants exclusively detected by the Hybrid pipeline.

**Figure 10.**
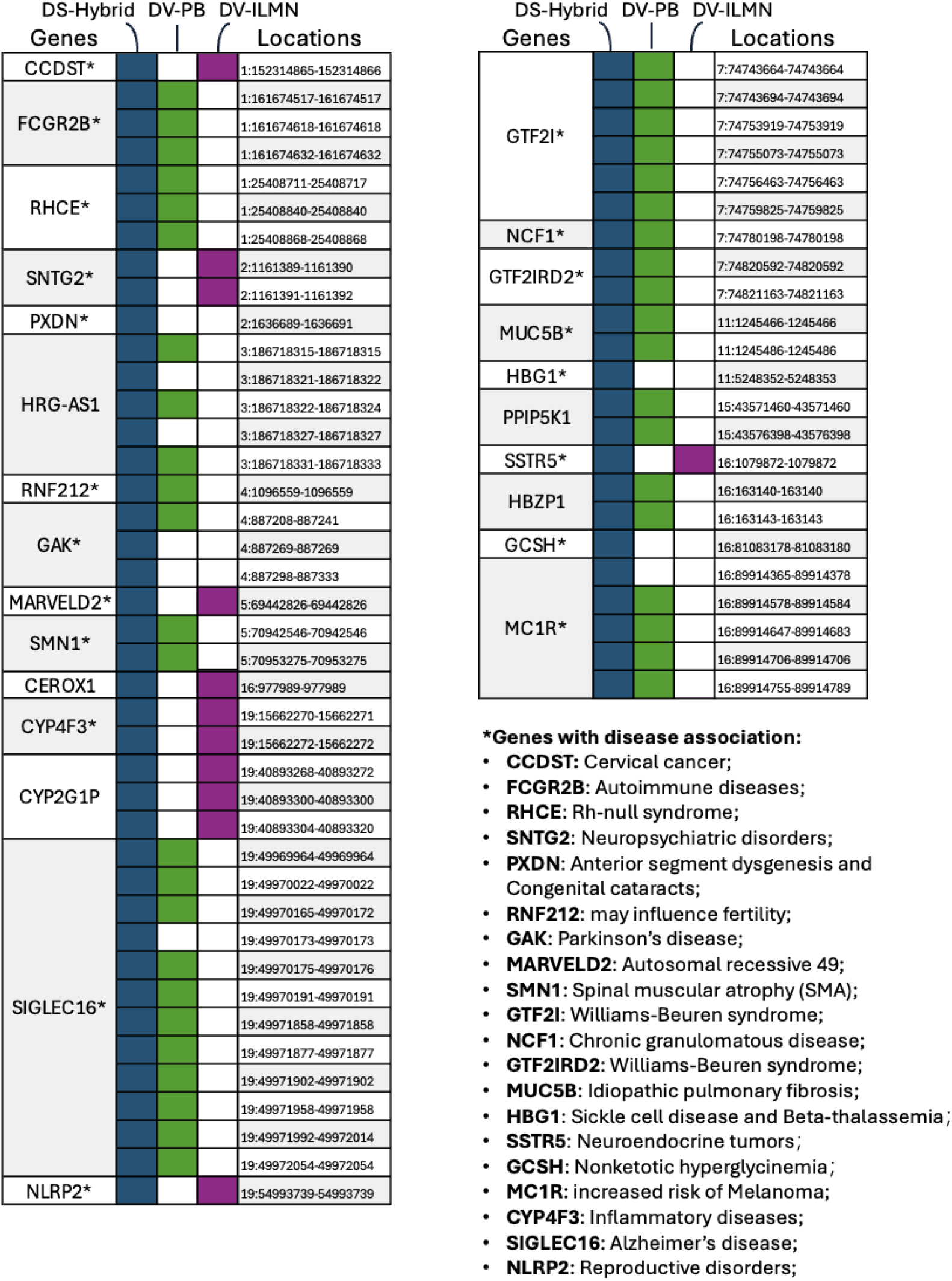
Identification landscape of selected SNPs/Indels missed by different pipelines. Variants were selected based on the intersection of HG002 CMRG exonic variants and those identified by one or two of the three pipelines. Many of the CMRG genes have well-documented disease associations, as shown in the right panel.

We also analyzed a previously published set of clinically relevant germline variants identified from 100 real patient samples^30^. Some variants in these patients were initially identified whole-exome sequencing, while variants other were identified using other molecular diagnostic approaches including Sanger sequencing and molecular ligation-based probe amplification (MLPA) as these variants are difficult to detect from traditional short-read sequencing. From this dataset, we selected all 42 SNP/Indel variants for further analysis. We generated simulated Illumina short-reads at 30x coverage and PacBio HiFi long-reads at 10x coverage and processed the simulated data through the DNAscope Hybrid pipeline. The DNAscope Hybrid pipeline successfully detected all 42 SNP/Indels. Among the identified variants, we focused on those previously reported in clinical cases, as shown in Table 1 below.

**Table 1.** Selected NGS challenging clinical SNP/Indels carried by patients diagnosed by traditional detection methods.

| Patient Sample ID | Case reported associated with this variant | Variant Details | Gene Locus | Original Diagnostic Method | Detected by DNAscope Hybrid |
| --- | --- | --- | --- | --- | --- |
| P3-E11 | Hearing Loss (RCV000151940.7) | Chr15(GRCh38):g.43600609_43600610delinsAG | STRC | Exome sequencing | True |
| P10-A4 | Congenital Adrenal Hyperplasia (CAH) | Chr6(GRCh38):g.32040535C>T | CYP21A2 | Sanger sequencing | True |
| P10-B4 | Congenital Adrenal Hyperplasia (CAH) | Chr6(GRCh38):g.32039081C>G | CYP21A2 | Sanger sequencing | True |
| P11-F11 | Nonsyndromic High Myopia and Reduced Cone Function | OPN1LW: LVAVA combination | OPN1LW; OPN1MW | Amplicon-based Long-read Sequencing | True |
| P50-D1 | Cone Dysfunction Syndromes | 1 copy OPN1LW: MIAVA combination | OPN1LW; OPN1MW | Molecular Ligation-based Probe Amplification (MLPA) + Sanger sequencing | True |
| P50-A5 | Hereditary Breast and Ovarian Cancer Syndrome (HBOC) | Chr17(GRCh38):g.43093577del | BRCA1 | single molecular Molecular Inversion Probes (smMIPs) | True |
| P50-B5 | Hereditary Breast and Ovarian Cancer Syndrome (HBOC) | Chr13(GRCh38):g.32333291dup | BRCA2 | single molecular Molecular Inversion Probes (smMIPs) | True |

A notable example is Patient P10-B4 (Figure 11 A), who was diagnosed with Congenital Adrenal Hyperplasia (CAH) caused by an SNP in intron 2 of the CYP21A2 gene. This variant disrupts normal splicing, leading to impaired 21-hydroxylase enzyme function. The CYP21A2 gene has a highly similar pseudogene (CYP21A1P), which creates challenges for short-read sequencing alignment due to their high sequence homology. Traditionally, Sanger sequencing was the only reliable method for detecting such variants. However, in this case, long-read assembled haplotype data was used to guide alignment, enabling accurate mapping of short reads and providing strong support for variant identification.

**Figure 11.**
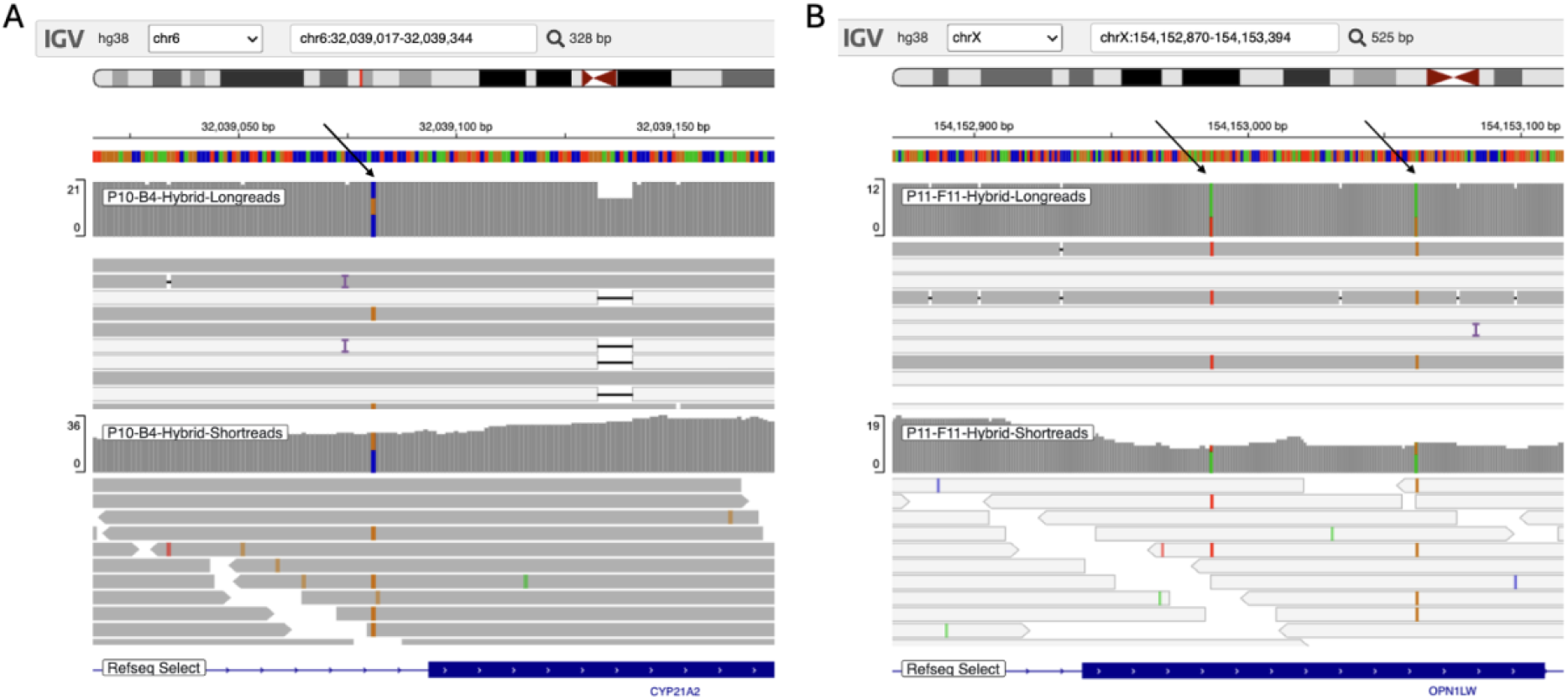
IGV screenshot showing variants: (A) Chr6(GRCh38):g.32039081C>G, carried by Patient P10- B4, and (B) OPN1LW: LVAVA combination, carried by Patient P11-F11. The upper tracks display the mapped results of simulated long reads, while the lower tracks show realigned short reads. Target variants are indicated by black arrows.

Figure 11-B shows simulated read data from patient P11-F11, who carries the LVAVA combination, which affects five key amino acid positions in exon 3 of the OPN1LW gene. OPN1LW plays a crucial role in the spectral tuning of the red-sensitive photopigment and mutations are associated with color vision impairment. The two critical DNA substitutions defining the LVAVA variant are highlighted by arrows, indicating their significance in modifying the opsin protein’s function. OPN1LW has a high degree of sequence identity to the other opsin genes, OPN1MW and OPN1SW, which creates challenges for short-read alignment across the opsin genes. The hybrid pipeline overcomes the difficulty in short-read alignment, providing sufficient coverage for accurate variant identification.

### Compute Resource Benchmark

A major challenge in whole genome sequencing secondary analysis is the long runtime, high cost of compute, and requirements for specialized hardware for obtaining an adequate TAT. The Sentieon software addresses these issue by running efficiently on commodity (x86 or Arm) CPU servers or workstations, offering accelerated runtimes, improved consistency, and high accuracy compared to other tools.

To assess the runtime of the Sentieon software, we tested three Sentieon pipelines – the DNAscope Hybrid pipeline with 10x PacBio HiFi and 35x Illumina data, DNAscope LongRead (PB) with 30x PacBio HiFi data, and DNAscope with 30x Illumina data. The benchmark assessed the runtime performance of alignment, preprocessing and SNP/Indel/SV/CNV calling. A 120 thread Azure instance (Standard HG120rs V3) was used as computation environment. The results for runtime, core-hours, and compute cost are shown in Table 2 below. The DNAscope LongRead and DNAscope pipeline runtime was previously published^31^.

**Table 2.**
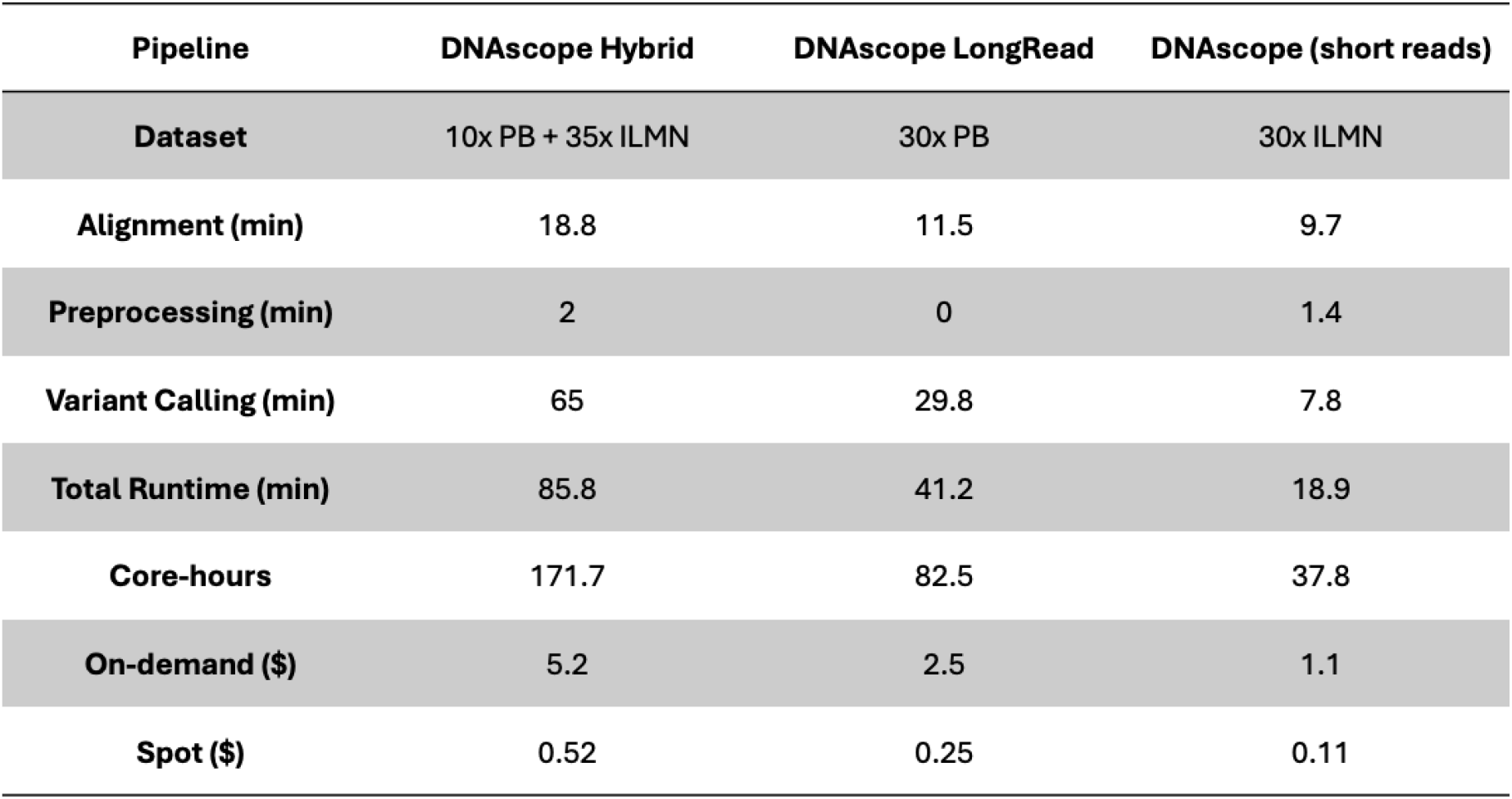
Compute resource benchmark for DNAscope pipelines. Benchmark environment is Azure Standard HB120rs v3 (120 vCPUs, 456 GiB memory, 512GB premium SSD), runtime and on-demand compute cost are displayed. The DNAscope Hybrid pipeline outputs SNP/Indel/SV/CNV, the DNAscope LongRead pipeline outputs SNP/Indel/SV, DNAscope short reads outputs SNP/Indel/CNV.

| Pipeline | DNAscope Hybrid | DNAscope LongRead | DNAscope (short reads) |
| --- | --- | --- | --- |
| Dataset | 10x PB + 35x ILMN | 30x PB | 30x ILMN |
| Alignment (min) | 18.8 | 11.5 | 9.7 |
| Preprocessing (min) | 2 | 0 | 1.4 |
| Variant Calling (min) | 65 | 29.8 | 7.8 |
| Total Runtime (min) | 85.8 | 41.2 | 18.9 |
| Core-hours | 171.7 | 82.5 | 37.8 |
| On-demand (\$) | 5.2 | 2.5 | 1.1 |
| Spot (\$) | 0.52 | 0.25 | 0.11 |

The DNAscope Hybrid pipeline is actively being developed, with future releases expected to show incremental improvements in computational efficiency and accuracy. Benchmarking results indicate that all three Sentieon pipelines completed the fastq-to-VCF analysis in approximately 20 minutes to less than 90 minutes for a cost of between $0.11 and $5.20 depending on data and spot or on-demand pricing.

## Methods

### Datasets used in this study

FASTQ files were downloaded from public source:

PacBio: Human whole genome sequencing datasets from the Revio system for the Genome in a Bottle trio HG002+HG003+HG004, with one Revio SMRT Cell per sample replicate^32^.

Illumina: pFDA Truth Challenge V2^33^.

Benchmark VCFs: SNP/Indel: NIST V4.2.1; draft Q100 V0.019; CMRG V1.00; SV: Q100 V0.019; CMRG V1.00

### DNAscope Hybrid Pipeline Overview

The Sentieon DNAscope Hybrid pipeline is designed to process and integrate both short and long sequencing reads from the same sample, achieving the most comprehensive and accurate variant calling results. This integrated approach ensures that variant calling accuracy surpasses the results obtained by processing short or long reads separately. The pipeline takes Fastq or BAM files as input and produces SNP, Indel, SV, and CNV calls in VCF format as output. It can be applied to any human whole genome sequencing assay and has potential for extension to other applications, such as whole exome sequencing, CMRG sequencing, or HLA analysis.

As shown in Figure 1, the DNAscope Hybrid pipeline aligns both short and long-reads to the reference genome using aligners specific for each data type (BWA-MEM for short-reads and minimap2 for long reads). At regions of the genome where short-read alignment is unreliable, the pipeline generates haplotypes from the long read alignments, and uses the long-read haplotypes to place the short-reads along the reference genome. In the final step, SNPs and Indels are called using all mapped reads and the called variants are genotyped and filtered using a machine learning model, similar to the Sentieon DNAscope pipeline^22^. The machine learning model used in the DNAscope Hybrid pipeline is trained using the GIAB v4.2.1 benchmark, with the HG003 sample held out from training. SVs are called using long reads only through the DNAscope LongRead SV module, and CNVs are called using short reads only with the CNVscope module. A single-line command interface (CLI) has been developed for users to easily run the pipeline by defining the input, output, and key parameters^34^.

sentieon-cli dnascope-hybrid [-h] \

-r REFERENCE \

--sr-aln SR_ALN [SR_ALN …] \

--lr_aln LR_ALN [LR_ALN …] \

-m MODEL_BUNDLE \ [-d DBSNP] \

[-b DIPLOID_BED] \

[-t NUMBER_THREADS] \

sample.vcf.gz

In addition to the DNAscope Hybrid pipeline, the following Sentieon pipelines and modules were utilized in this benchmark: BWA/MM2 alignment (which supports machine learning models), DNAscope LongRead, DNAscope LongRead SV, CNVscope, and Hap-eval.

### Sentieon Alignment

Sentieon released an accelerated version of BWA-MEM^35,36^ in 2017^37^ and followed with an accelerated version of Minimap2^38^ in the 202010.04 release. These tools provide results consistent with the open-source versions but deliver 2-3x faster performance. In the 202308 release, a new version of these two alignment tools was introduced, improving whole genome sequencing (WGS) alignment runtimes by approximately 2x by using a model file to optimize compute resources.

For this benchmark, Illumina NovaSeq6000 reads from HG002 samples and other two GIAB samples (downloaded from the precisionFDA Truth V2 challenge) were mapped to the GRCh38 reference genome (GCA_000001405.15_GRCh38_no_alt_analysis_set_maskedGRC_exclusions_v2.fasta). Reads from PacBio Revio were mapped to the GRCh38 reference genome using Sentieon MM2.

### Short Variant Calling

In addition to the DNAscope Hybrid pipeline, down-sampled long-read datasets were also analyzed using the DNAscope LongRead pipeline^23^. Originally published in collaboration with PacBio in 2022, DNAscope LongRead performs mapping, phasing, and utilizes pre-trained machine learning models to correct sequencer-specific error patterns and accurately call short variants. Subsequent releases added support for ONT reads, further enhancing pipeline performance. The DNAscope Longread pipeline can also be run via Sentieon CLI, with the following command line:

sentieon-cli dnascope-longread [-h] \

-r REFERENCE \

--fastq INPUT_FASTQ … \

--readgroups READGROUP … \

-m MODEL_BUNDLE \ [-d DBSNP] \

[-b DIPLOID_BED] \

[-t NUMBER_THREADS] \

[-g] \

--tech HiFi|ONT \

[--haploid-bed HAPLOID_BED] \

sample.vcf.gz

DeepVariant (v1.8.0)^28^ was used to generate variants from long-read only data, using its default settings. Dragen accuracy metrics were generated from VCF files downloaded from a recently published work^39^.

Benchmarking of SNV and Indels was conducted using hap.py with the GIAB V4.2.1, draft Q100, and CMRG benchmark VCFs and BED files.

### Structural Variant Calling and Benchmarking with Hap-eval

DNAscope LongRead SV caller integrated in the DNAscope Hybrid pipeline. It performs haplotype-resolved SV calling and works well with both PacBio HiFi and ONT data, supporting various sequencing chemistry versions and base callers. Pbsv (v2.10.0)^40^ was also used to generate long-read-only SVs. Dragen v4.2 SVs were obtained from an earlier publication^40^.

Benchmark of structural variants (SV) was performed using “hap-eval,”^41^ an open-source structural variant benchmarking tool developed by the Sentieon team. Sentieon developed hap-eval to address some limitations of the popular tool “Truvari,”^42^. Older versions of Truvari performed a pairwise comparison of variants, without accommodation for multiple nearby variants or multi-allelic sites. Hap-eval assembles multiple variants into haplotypes and conducts haplotype-based comparisons. This VCF comparison engine is assembly-based and compares SV haplotypes at a single-base resolution. Hap-eval works in four steps: 1) Combine base and comparison calls in sorted order; 2) Create comparison chunks; 3) Perform a 1-to-1 comparison within each chunk between base and comparison calls to create a match matrix. 4) Make calls based on the matching matrix.

### CNV Calling

In DNAscope Hybrid, CNV identification is performed by CNVscope, a short-read WGS CNV caller released in Sentieon version 202308.03. CNVscope is designed for germline CNV calling across diploid chromosomes, identifying events greater than 1kb in length from a 30x WGS dataset. The CNVscope algorithm uses read-depth profiling, normalization, feature collection, and segmentation to identify CNV events. Identified events are then filtered using a pre-trained machine learning model. The model was trained and tested using datasets developed from the HPRC assemblies and T2T assemblies, including the Q100 assembly. CNVscope uses a reference independent approach and can be used with hg38, b37, or other high quality reference genome assemblies of diploid organisms.

sentieon driver -t NUMBER_THREADS -r REFERENCE -i DEDUPED_BAM **\**

--algo CNVscope --model ML_MODEL/cnv.model TMP_VARIANT_VCF

sentieon driver -t NUMBER_THREADS -r REFERENCE --algo CNVModelApply **\**

--model ML_MODEL/cnv.model -v TMP_VARIANT_VCF VARIANT_VCF

We also generated CNV calls using CNVnator (v0.4.1)^43^. Benchmarking of CNVs was conducted using an in-house developed script. The evaluation is between the call and truth considers a call to be TP is it has at least 30% overlap compared to both the truth and call intervals, and they need both either gain or loss. The expanded truth interval is considered here if the CNV event occurs at a segmental duplication region.

CNV benchmark for HG002 was generated based on draft Q100 SV benchmark, where long indel are converted to CNV events. In the conversion from INDEL to CNV events, for long deletion, the CNV loss event is where the deletion occurs, and the copy number state is determined by the genotype. For long insertions, the inserted sequence is matched locally and throughout the genome to identify matches. If a match is found, a CNV gain event is found. For both insertion and deletion, it will also look for repeats within the indel sequence and around the CNV event intervals, and the CNV event interval may be expanded if needed.

## Discussion

Here we present the DNAscope Hybrid pipeline, a robust, fast and accurate pipeline for combined short and long-read data. The DNAscope Hybrid pipeline uses a novel approach, with long-read haplotypes guiding short-read alignment. We demonstrate that this approach enables higher variant calling accuracy than single-technology pipelines. In addition to SNVs/indels, the hybrid pipeline incorporates specialized callers for structural and copy-number variation to enable accurate detection of all types of variation.

Variant calls from the DNAscope Hybrid pipeline and other tools were benchmarked using multiple samples and multiple benchmark datasets. We utilized three samples (HG002, HG003, and HG004) from the GIAB v4.2.1 benchmark to confirm that our method works well across multiple samples and is not overfit to the samples used in model training (HG003 is held out during the training process). We also tested the performance of these tools using the CMRG benchmark and the draft Q100 benchmark. The CMRG benchmark contains genomic regions that are difficult to resolve with short-read sequencing, while the draft Q100 benchmark uses an assembly-based approach to extend into regions that are difficult to resolve with traditional mapping-based approaches. Taken together, these benchmarks validate the performance of the DNAscope Hybrid pipeline as a robust pipeline for the analysis of combined short and long-read data.

Sequence coverage is a major consideration during project planning, for both clinical and research projects. The DNAscope Hybrid pipeline can be used with a range of coverages, including targeted long-read sequencing, and will attempt to fully utilize the available read data. In this paper we benchmark full-coverage short-read datasets (at ∼35x coverage) with a range of long-read coverages to assess pipeline performance. Given the robust performance of the hybrid pipeline at a range of coverages, we believe full-coverage short-read sequencing combined with either targeted or low-coverage (7x to 15x) long-read sequencing will be applicable to a wide-range of projects; enabling high accuracy SNVs and Indels while also incorporating high accuracy structural variants, which are not accessible from short-read data alone.

### Future Directions

The DNAscope Hybrid Pipeline is actively under development, with several planned improvements and expansions.

Somatic variant calling is an area where the hybrid method could provide significant advantages. Clinically relevant somatic variants often occur at low allele frequencies and may introduce homopolymer sequence repeats due to deficiencies in DNA error repair mechanisms. Somatic structural variants, including gene fusions, are important drivers of tumorigenesis. Accurately detecting these variants requires both high sequencing depth and long reads to resolve complex genomic regions. Currently, no single sequencing technology meets both requirements effectively. Hybrid approaches present a promising solution by combining the strengths of short- and long-read sequencing technologies, especially when combined with targeted sequencing of clinically relevant regions that are difficult to resolve from short-read data alone.

In addition to single-sample variant identification, DNAscope can process cohort samples through joint calling, utilizing pre-trained sequencing platform-specific models for short- or long-read data. This approach represents an alternative method of hybrid analysis across multiple sample types. In a recently published application note^44^, the DNAscope joint caller demonstrated its ability to harmonize different error patterns across multiple sequencing platforms, producing results that are fully interoperable with datasets generated from Illumina sequencers. Moving forward, we plan to collect additional short- and long-read datasets from the 1kGP project and further benchmark the hybrid joint calling pipeline using these data.

Currently, this benchmark project includes datasets from only two sequencing platforms: Illumina and PacBio. However, several other commercially available platforms—particularly for short reads—are gaining traction. DNAscope already supports additional sequencing platforms, including Element Biosciences, Ultima Genomics, Complete Genomics, among others. We anticipate that the hybrid pipeline will also be applicable to these and other emerging sequencing platforms. As part of our roadmap, we plan to develop and benchmark hybrid models for these additional sequencing platforms.

## Supplementary Datasheets

Table S1-S10 contains datasheets and more detailed information for figures in the main manuscript.

## Data Availability

**All VCF files and CNV truth files have been uploaded to** https://zenodo.org/records/15033928

## Code Availability

Scripts used in this study can be found in https://github.com/Sentieon/sentieon-cli. Sentieon software package is freely available for trial purpose upon request.

## Completing Interests

All authors are employees of Sentieon Inc. and own stock options as part of their standard compensation package.

